# Host lung environment limits *Aspergillus fumigatus* germination through a SskA-dependent signaling response

**DOI:** 10.1101/2021.08.27.456493

**Authors:** Marina E. Kirkland, McKenzie Stannard, Caitlin H. Kowalski, Dallas Mould, Alayna Caffrey-Carr, Rachel M. Temple, Brandon S. Ross, Lotus A. Lofgren, Jason E. Stajich, Robert A. Cramer, Joshua J. Obar

## Abstract

*Aspergillus fumigatus* isolates display significant heterogeneity in growth, virulence, pathology, and inflammatory potential in multiple murine models of invasive aspergillosis. Previous studies have linked the initial germination of a fungal isolate in the airways to the inflammatory and pathological potential; but the mechanism(s) regulating *A. fumigatus* germination in the airways are unresolved. To explore the genetic basis for divergent germination phenotypes, we utilized a serial passaging strategy in which we cultured a slow germinating strain (AF293) in a murine lung based medium for multiple generations. Through this serial passaging approach, a strain emerged with an increased germination rate that induces more inflammation than the parental strain (herein named <u>L</u>ung <u>H</u>omogenate <u>Evol</u>ved (LH-EVOL)). We identified a potential loss of function allele of *Afu5g08390* (*sskA*) in the LH-EVOL strain. The LH-EVOL strain had a decreased ability to induce the SakA-dependent stress pathway, similar to AF293 Δ*sskA* and CEA10. In support of the whole genome variant analyses, *sskA*, *sakA*, or *mpkC* loss of function strains in the AF293 parental strain increased germination both *in vitro* and *in vivo*. Since the airway surface liquid of the lungs contains low glucose levels, the relationship of low glucose concentration on germination of these mutant AF293 strains was examined; interestingly, in low glucose conditions the *sakA* pathway mutants exhibited an enhanced germination rate. In conclusion, *A. fumigatus* germination in the airways is regulated by SskA through the SakA MAPK pathway and drives enhanced disease initiation and inflammation in the lungs.

**IMPORTANCE:** *Aspergillus fumigatus* is an important human fungal pathogen particularly in immunocompromised individuals. Initiation of growth by *A. fumigatus* in the lung is important for its pathogenicity in murine models. However, our understanding of what regulates fungal germination in the lung environment is lacking. Through a serial passage experiment using lung-based medium, we identified a new strain of *A. fumigatus* which has increased germination potential and inflammation in the lungs. Using this serially passaged strain we found it had a decreased ability to mediate signaling through the osmotic stress response pathway. This finding was confirmed using genetic null mutants demonstrating that the osmotic stress response pathway is critical for regulating growth in the murine lungs. Our results contribute to the understanding of *A. fumigatus* adaptation and growth in the host lung environment.

## INTRODUCTION

*Aspergillus fumigatus* is a ubiquitous mold essential for decomposition of organic material that can easily become airborne. Infection is established in the respiratory tract by inhalation of dormant conidia which are roughly 2 μm in diameter. Their small size facilitates embedment into tissues if those resting conidia are not removed through mucociliary action of respiratory epithelial cells. If allowed sufficient time, *A. fumigatus* conidia will begin to germinate, which initiates with osmotic swelling and then polarized growth of germ tube. At this point, a healthy immune system can detect pathogen associated molecular patterns (PAMPs) on the cell surface of swollen or germinated conidia, clearing the swollen conidia before disease can be established. Should this occur in an immunocompromised host, germination can go unchecked resulting in fungal biofilm formation and tissue invasion establishing severe disease^1–3^.

Previous work has shown a correlation between germination and pathogenicity of *A. fumigatus* in a strain-dependent manner in a model of bronchopneumonia^4^. Between clinical isolates AF293 and CEA10, CEA10 exhibits faster germination both *in vitro* and *in vivo* in murine lung airways^4^, as well as in a zebrafish infection model^5^. This increased germination corresponds with greater tissue damage and greater inflammation following CEA10 challenge compared to AF293 challenge^4–6^. However, conditions and genetic pathways that regulate the germination rate for each strain in the lung microenvironment remain unresolved.

For germination to occur conidia must first sense whether their environment has sufficient nutrient availability and the presence of potential stressors. In *Aspergillus nidulans*, the presence of glucose was sufficient to induce germination of conidia^7^, while in *A. fumigatus* glucose in the presence of water and oxygen to drive cellular respiration was necessary for conidia germination^8^. The germination process is regulated by both Ras signaling^9^ and G-protein signaling through the cAMP-PKA signaling pathway^8–10^. Additionally, calcium signaling through the calcineurin pathway and the CrzA transcription factor is needed for *A. fumigatus* germination^11,12^. Further, in *A. fumigatus* nitrogen sources, such as ammonium chloride or proline, drive more robust germination in a SakA-regulated manner^13^. Once conidia begin to germinate, they begin an isotropic growth phase which modifies the composition of their cell walls, subjecting conidia to detection by a host’s immune system. Polarized growth follows as swollen conidia develop into germlings then hyphae before establishing a complex network called mycelia^1^.

The lung itself has evolved to interact constantly with an onslaught of foreign particulate matter and microbes in the air. Lung airways are a low-nutrient environment: airway surface liquid (ASL) glucose concentrations in a healthy adult are approximately 0.4 m*M*, which is about 10-fold less than in the plasma^14^. Reduced nutrient availability is thought to limit microbial growth in the lung environment. In addition, the ASL facilitates the first line defense of airway epithelial cells by providing mucous and water secretions which assist in mucociliary clearance^14–16^. The ASL can become dysregulated in chronic disease states brought on by asthma^17,18^, COPD^18^, cystic fibrosis^19^, and type II diabetes^19–21^ resulting in increased glucose levels^21^, all of which have increased risk for developing aspergillosis ^22–25^.

In this study, we explored the molecular mechanisms regulating the germination of *A. fumigatus* in context of the murine lung environment. To do this we utilized a serial passaging approach by culturing the slow germinating AF293 strain in lung homogenate medium and selecting for increased germination after multiple generations. A novel AF293-dervived strain, LH-EVOL, was selected and found to germinate quicker in the murine lung airways compared to the parental AF293 strain resulting in a stronger proinflammatory IL-1α-dependent immune response. Whole genome sequencing of LH-EVOL revealed conserved mutations in *Afu5g08390* (*sskA*), which is part of the SakA MAPK stress pathway^26^. *In vitro* and *in vivo* growth kinetics of an AF293 Δ*sskA* mutant recapitulated the quick germinating phenotype of LH-EVOL. Interestingly, this rapid germination phenotype was further accelerated *in vitro* under low glucose conditions, which more closely resemble the concentration of glucose found in the lung environment. Taken together, this study further supports previous observations that germination rate is a key step in determining the qualitative and quantitative inflammatory immune response to *A. fumigatus* strains in the murine lung and identifies the SakA-pathway as a key mediator of host lung environment.

## MATERIALS AND METHODS

### Fungal strains and growth conditions

All fungal strains used in this study are listed in Supplemental Table 1. For conidial harvest, strains were cultured on 1% glucose minimal media (GMM; 1% or 55.56 m*M* glucose, 6 g/L NaNO_3_, 0.52 g/L KCl, 0.52 g/L MgSO_4_•7H_2_O, 1.52 g/L KH_2_PO_4_ monobasic, 2.2 mg/L ZnSO_4_•7H_2_O, 1.1 mg/L H_3_BO_3_, 0.5 mg/L MnCl_2_•4H_2_O, 0.5 mg/L FeSO_4_•7H_2_O, 0.16 mg/L CoCl_2_•5H_2_O, 0.16 mg/L CuSO_4_•5H_2_O, 0.11 mg/L (NH_4_)_6_Mo_7_O_24_•4H_2_O, and 5 mg/L Na_4_EDTA; pH 6.5) agar plates for three days at 37°C, at which time conidia were collected by flooding plates with 0.01% PBS/Tween 80 and gently scraping mycelia using a cell scraper. Conidia were then filtered through sterile Miracloth, washed, and resuspended in phosphate buffered saline (PBS), and counted on a hemacytometer. Strains were stored in PBS and were used immediately for *in vitro* experiments or allowed to rest overnight for *in vivo* studies.

### Serial passaging of AF293 through murine lung homogenate medium

The wild-type AF293 strain of *A. fumigatus* was used for a serial passage experiment as it germinates slowly in the lung airways^6^. Murine lung homogenate medium was made as previously described^6^, where naïve C57BL/6J lungs were crushed in 2 ml PBS and filtered through a 70 μM filter after which cellular debris was removed by centrifugation. The working concentration used was 1:4 in PBS and contains ~35μM glucose, which was determined using a glucose oxidase assay kit. AF293 was cultured in lung homogenate medium for 8 hours at 37°C shaking at 350 rpm to induce germination. The fungal material was then collected and plated on 1% GMM agar plates and grown for 3 days at 37°C. The population of conidia were then collected and cataloged. This process was repeated 13 times. LH-EVOL was then grown under standard growth conditions prior to further analyses. Twenty individual clones of the LH-EVOL population were isolated to a single spore for further analysis in our lung homogenate germination assay.

### Whole genome sequencing and variant identification

To assess potential mutations within the LH-EVOL strain whole genome DNA sequencing was performed on three LH-EVOL clones (LH-EVOL clone 4, LH-EVOL clone 11, and LH-EVOL clone 12) with enhanced germination in lung homogenate medium. Genomic DNA was extracted from mycelia grown in Petri plates using liquid 1% GMM supplemented with 0.5% yeast extract, following to previously published methods^27^. The DNA concentration was quantified using a Qubit 2.0 Fluorometer (Invitrogen) and the manufacturer’s recommended Broad Range protocol. The genomic DNA sequencing was genomic sequencing libraries were prepared by SeqMatic (Fremont, CA) using Illumina TruSeq DNA kits and sequenced on MiSeq Illumina sequencer in 2×250 bp format following manufacturer recommendations for paired end library construction and barcoding. The sequencing produced a range of 1 - 1 average of 1.1M – 1.6M reads, which is from 300-440 Mb and 10-14X coverage of the *A. fumigatus* genome.

The DNA sequence reads for each strain were aligned to the AF293 reference genome downloaded from FungiDB v.39^28,29^ with BWA v0.7.17^30^ and converted to BAM files using SAMtools v1.10^31^. Following best practices^32^, the reads were marked for PCR and optical duplication using picard tools v2.18.3 (http://broadinstitute.github.io/picard). For variants identified near inferred INDELs, reads were realigned using RealignerTargetCreator and IndelRealigner in the Genome Analysis Toolkit GATK v3.7^32^. The SNPs and INDELS were genotyped relative to AF293 using HaplotypeCaller in GATK v4.1.1.0^33^. Filtering was accomplished with GATK’s SelectVariants with the following parameters: for SNPS: -window-size = 10, -QualByDept < 2.0, -MapQual < 40.0, -QScore < 100, -MapQualityRankSum < −12.5, -StrandOddsRatio > 3.0, -FisherStrandBias > 60.0, -ReadPosRankSum < −8.0. For INDELS: -window-size = 10, -QualByDepth< 2.0, -MapQualityRankSum < −12.5, -StrandOddsRatio > 4.0, -FisherStrandBias > 200.0, -ReadPosRank < −20.0, -InbreedingCoeff < −0.8. The genic overlap and consequence of the variants were classified with snpEff^34^. Variants were transformed into tabular format from VCF format using script ‘snpEff_2_tab.py’.

The genomic short reads for the four strains are deposited in NCBI SRA associated with BioProject PRJNA757188. Scripts for running the pipeline for analyses are available in the github (https://github.com/stajichlab/Afum_LH-EVOL).

### *Aspergillus fumigatus* pulmonary challenge model

C57BL/6J mice (Stock #000664) were purchased from Jackson Laboratory at 7-8 weeks of age. *Il1a^(−/−)^* mice^35^ were bred in-house at Dartmouth College. Animal studies were carried out in accordance with the recommendations in the Guide for the Care and Use of Laboratory Animals. The animal experimental protocol 00002168 was approved by the Institutional Animal Care and Use Committee (IACUC) at Dartmouth College. Mice were challenged with 4×10^7^ conidia in PBS intratracheally under isoflurane anesthesia in a volume of 100μl. At the indicated experimental time points mice were euthanized and bronchoalveolar lavage fluid (BALF) was collected by washing the lungs with 2 ml of PBS containing 0.05 *M* EDTA. BALF was clarified by centrifugation and stored at −20°C until analysis. After centrifugation, the cellular component of the BAL was resuspended in 200 μl of PBS. BAL cells were subsequently spun onto glass slides using a Cytospin4 cytocentrifuge (Thermo Scientific) and stained with Diff-Quik stain set (Siemens) for quantification of fungal germination, as previously described^4^. The percent germination of each *A. fumigatus* strain was quantified by manual counting of 100-200 fungal conidia and germlings at 100X magnification using a standard upright microscope. For histological analysis lungs were filled with and stored in 10% buffered formalin phosphate for at least 24 hours. Lungs were then embedded in paraffin and lung sections were stained with Grocott’s methenamine silver (GMS) staining to assess fungal germination.

### Murine lung RNA preparation and NanoString analysis

Animals were challenged with either PBS, AF293, or LH-EVOL, as described above. Forty hours after inoculation mice were euthanized and whole lungs were removed for mRNA analysis. Lungs were placed in 2 ml of TRIzol and homogenized with a glass dounce homogenizer, followed by treatment with chloroform to extract RNA according to manufacturer’s instructions. After RNA was assessed for quality, 100 ng of RNA was used per reaction using the nCounter Mouse PanCancer Immune Profiling Panel (NanoString). nSolver 4.0 software was used for background subtraction and normalization. nSolver Advanced Analysis 2.0 was used for quality control analysis and pathway analysis.

### Mutant *Aspergillus fumigatus* strain construction

The Δ*sskA* strain was generated in the uracil/uridine auxotrophic strain AF293.1 through homologous recombination. The gene replacement construct was generated using overlap extension PCR as previously described^36^ in which ~1 kb of sequence 5’ to the start codon of *sskA* and ~1 kb 3’ to stop codon of *sskA* were amplified and fused to *Aspergillus parasiticus* orotidine 5’-monophosphate decarboxylase gene, *pyrG.* This *pyrG* fragment was amplified from pSD38.1 where *pyrG* has been blunt-end ligated into pJET vector (Thermo Scientific, #K1231) with 5’ and 3’ linker sequences, 5’-accggtcgcctcaaacaatgctct-3’ and 5’-cgcatcagtgcctcctctcagac-3’, respectively^37^. Primers for the amplification of this gene replacement construct are provided in Supplemental Table 2. The transformation of AF293.1 to generate Δ*sskA* was carried out as previously described for the transformation of *A. fumigatus* protoplasts generated using Lysing Enzyme from *Trichoderma harzianum* (Sigma: L1412)^38^. Candidate Δ*sskA* strains were selected for restored prototrophy. Single spores were isolated from the candidate strains to generate clonal strains that were confirmed to be Δ*sskA* through PCR and Southern Blot analyses. Southern blot analyses were performed as previously described using the digoxigenin-anti-digoxigenin detection system (Roche Diagnostics)^38^.

For the generation of the *sskA^RC^* construct, the pBluescript II KS(+) (Addgene) was utilized. The multiple cloning site was expanded and the pyrithiamine resistance cassette was amplified from pPTR I (Takara) to include XhoI (New England Biolabs) restriction sites at the 5’ and 3’ ends and introduced via the SalI site (pSD11). From AF293 genomic DNA we amplified a ~4 kb fragment that included ~1kb 5’ and ~200 bp 3’ of *sskA* and introduced PacI and NotI (New England Biolabs) restriction sites at the 5’ and 3’ ends, respectively. The *sskA^RC^* construct, pSskA-R was confirmed through restriction digestion and Sanger sequencing. The construct was isolated (Zyppy Plasmid Miniprep, Zymo Research) and transformed into the Δ*sskA* fungal genome as described above. The *sskA^RC^* strain, which contained an ectopic insertion of the *sskA* wild-type allele, was confirmed with qRT-PCR as previously described^37^. The housekeeping genes utilized were *tub2* and *actA.*

### Fungal germination assays

*In vitro* fungal germination assays were conducted as previously described^6^. Briefly, fungal strains were inoculated in either 1% GMM (55.56 mM of glucose), 0.25% GMM (13.89 mM of glucose), 1% GMM supplemented with 0.5% yeast extract (YE), or lung homogenate at a concentration of 1×10^7^conidia/ml in 2 ml of medium in glass 20ml disposable scintillation vials (VWR). Cultures were incubated at 37°C while shaking at 300 rpm. Starting at 4 hours a wet mount was made every hour to count the number of germlings to conidia. Samples were vortexed with 1.0 mm beads to break up clumps before mounting and fungal germination was quantified manually using the 40× objective lens of an upright VWR microscope. A minimum of 100 conidia and germlings for each sample were counted.

### Radial growth assay for testing response to osmotic stress

GMM plates (1% glucose) were supplemented with 1M NaCl, 1M KCl, 0.5M CaCl_2_, or 1M Sorbitol. 1×10^5^ conidia were inoculated onto plates in 2 μl drops. Plates were incubated at 37°C for 72 hours, at which point the diameter of the colony was measured. Inhibition was determined by normalizing each strain to their 72 hour GMM growth.

### Fungal RNA extraction and qRT-PCR analysis for SakA-dependent transcripts

For transcript analysis during growth during osmotic stress, 1×10^6^ conidia of AF293, AF293 Δ*sakA*, and LH-EVOL were grown in a 6-well plate in 5 mL of GMM media for 30 hours at 37°C. Mycelia were transferred to a new 6-well plate with either GMM or GMM supplemented with 1M NaCl under sterile conditions and incubated for 30 minutes at 37°C. Mycelia were transferred to ZR BashingBead Lysis Tubes (0.1 & 0.5 mm, Zymo Research) and homogenized at 4°C using a bead beater for 4 minutes at 250 RPM. RNA was purified from lysate following the manufacturer’s protocol for Quick-RNA Fungal/Bacterial Miniprep Kit (Zymo Research). cDNA was synthesized following the manufacturer’s protocol for QuantiTect Reverse Transcription Kit (Qiagen) including gDNA removal. Quantitative PCR was performed using SsoAdvanced Universal SYBER Green Supermix according to manufacturer’s directions and primers described in Supplemental Table 3 using the thermocycler Bio-Rad C-1000 CFX96. Cycling parameters were according to SsoAdvanced Universal SYBER Green Supermix manufacturer recommendation using 60°C as the annealing temperature. Transcripts were normalized to *actA* using the 2^−ΔΔCt^ method^39^.

Protein Sequence Analysis. Comparison of proteins of the SakA signaling network between CEA10 and Af293 was done using the web Protein BLAST (BLASTp) platform from the US National Library of Medicine, National Center for Biotechnology Information (https://blast.ncbi.nlm.nih.gov/Blast.cgi) using the non-redundant protein sequence database. Protein sequences used for this analysis are listed in Supplemental Table 4.

### Statistical analysis

All graphs and statistical analyses were conducted using the GraphPad Prism 9 software. Statistical significance between experimental groups was determined using Student’s t-test (comparison of two experimental groups normally distributed), a Mann-Whitney U test (comparison of two experimental groups that are not normally distributed), a one-way ANOVA using Dunn’s post-test (comparison of more than two experimental groups), or two-way ANOVA using Tukey’s post-test (comparison of more than two experimental groups with two variables) or Kruskal-Wallis post-test for non-gaussian distributions.

## RESULTS

### *In vitro* serial passaging of AF293 in lung homogenate medium leads to a strain with increased airway growth

Previously, we had established that strains of *Aspergillus fumigatus*, in the context of the lung environment, can be grouped into fast and slow germinators who induce markedly different pathologies^4,40^. Notably, fast germinating strains, such as CEA10, induce extensive loss of alveoli and destruction of the parenchyma with substantial hemorrhaging when compared to slow germinating strains, such as AF293. To identify the potential molecular pathways responsible for the quick germination phenotype within the lung environment by *A. fumigatus* we utilized an experimental serial passage approach, which enables us to identify potential fungal regulators of growth in the lung microenvironment without knowing the exact nutrients that are biologically available. Specifically, the slow germinating AF293 strain was serially cultured in lung homogenate medium for 8h to initiate germination then plated on 1% GMM plates for 3 days before collecting spores for cataloging and the next passage (Figure 1A). After 13 passages we identified a novel AF293-based strain, referred to as LH-EVOL, that was able to germinate rapidly in lung homogenate medium (Figure 1B). The germination kinetics of the LH-EVOL strain in lung homogenate medium was significantly faster than its parental AF293 strain, while having similar germination kinetics to the CEA10 strain in this *in vitro* assay (Figure 1B). To determine if the LH-EVOL strain also has an increased germination rate during initial airway challenge of mice, C57BL/6J mice were inoculated with 4×10^7^ conidia of either parental strain AF293 or LH-EVOL. After challenge, significantly more germination by LH-EVOL within the BALF 12 hours post inoculation was observed compared to its parental AF293 strain (Figure 1C). Thus, our *in vitro* serial passaging experimental conditions drove the development of a novel AF293-based strain that can rapidly germinate within the lung airways of mice.

**Figure 1.**
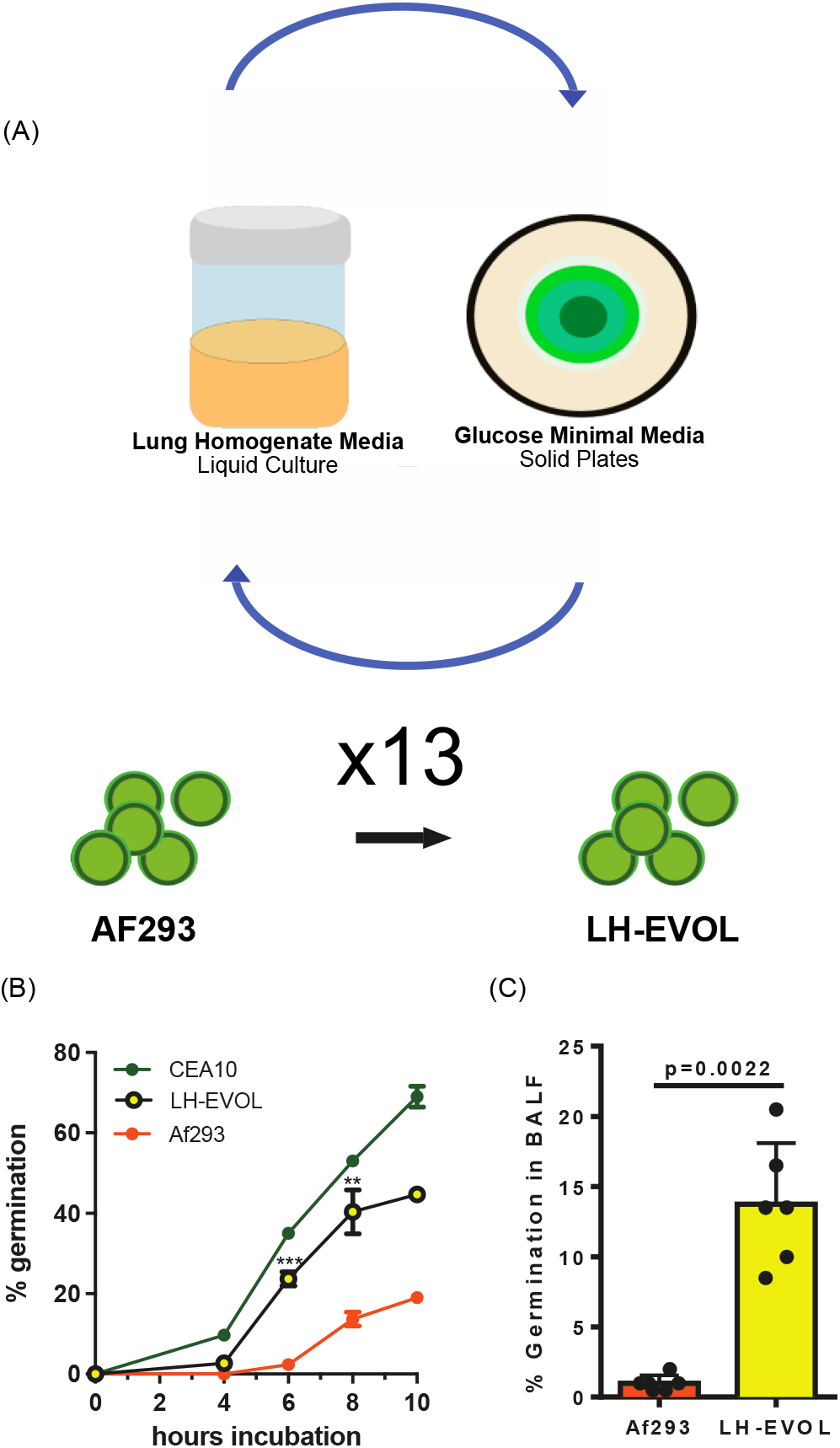
Lung evolved (LH-EVOL) *A. fumigatus* demonstrates increased *in vitro* and *in vivo* germination kinetics. (A) LH-EVOL was generated through serial passaging of the AF293 strain thirteen times in lung homogenate media. (B) Lung tissue from C57BL/6 mice was homogenized in PBS and used as the medium for *in vitro* germination assays. Percent germination over time in lung homogenate media of AF293; CEA10, or LH-EVOL. was quantified every 2 h by microscopically counting the numbers of conidia and germlings. Data are represented as the percentage of fungal matter that was germinated. Data are representative of at least 2 independent experiments consisting of 3 biological replicates per group. Each symbol represents the group mean ± 1 standard error of the mean. (C) C57BL/6J mice were challenged with 4×10^7^ conidia of AF293 or LH-EVOL. At 12 hpi, mice were euthanized, and BALF was collected to quantify germination in the airways. Data are representative of results from 2 independent experiments with 5 mice per group. (B, C) Statistical significance was assessed between LH-EVOL and AF293 using a Mann-Whitney U-test. (**) indicates a p-value ≤ 0.01; (***) indicates a p-value ≤ 0.001.

### Comparison of pulmonary inflammatory response induced following AF293 and LH-EVOL challenge in the murine lung

Given the different germination rates of the AF293 and LH-EVOL strains *in vivo*, we next analyzed the inflammatory response induced in the lung after challenge with each respective strain. Total RNA was extracted from whole lungs of C57BL/6J mice challenged 40 hours prior with either AF293 or LH-EVOL. Changes in mRNA levels of immune-related genes were analyzed using NanoString nCounter technology. We observed strikingly different overall changes in mRNA levels in PBS-challenged animals compared to the two fungal strains (Supplemental Dataset 1-2). Importantly, mRNA levels of 57 genes were significantly increased greater than 2-fold following LH-EVOL challenge when compared with AF293 challenge, whereas 65 genes were significantly increased greater than 2-fold following AF293 challenge when compared with LH-EVOL challenge (Figure 2A and Supplemental Dataset 3). When we examined differential gene expression and pathway analysis using nSolver Advanced Analysis software, we noted a dramatic signature related to IFN signaling following AF293 challenge (Figure 2B). This increased IFN signaling signature is evidenced by significant increases in *Cxcl9*, *Cxcl10, Stat1, Zbp1, Irf7, Isg15*, and *Ifit3* levels (Table 1). It has been established that early anti-fungal immune responses to conidia will induce expression of IFN-β and CXCL10^41^, therefore increased IFN signaling may occur in AF293 due to slower germination. In contrast, there were dramatic increases across several pathways associated with innate immunity and cytokine/chemokine signaling following LH-EVOL challenge relative to AF293 challenge (Figure 2B). Increased mRNA expression levels of pro-inflammatory cytokines including *Il1a, Il6, Csf2, Cxcl1, Cxcl2*, and *Cxcl5* and expression levels of the transcription factors *Nfkb1* and *Hif1a* were elevated following LH-EVOL challenge (Table 2).

**Figure 2.**
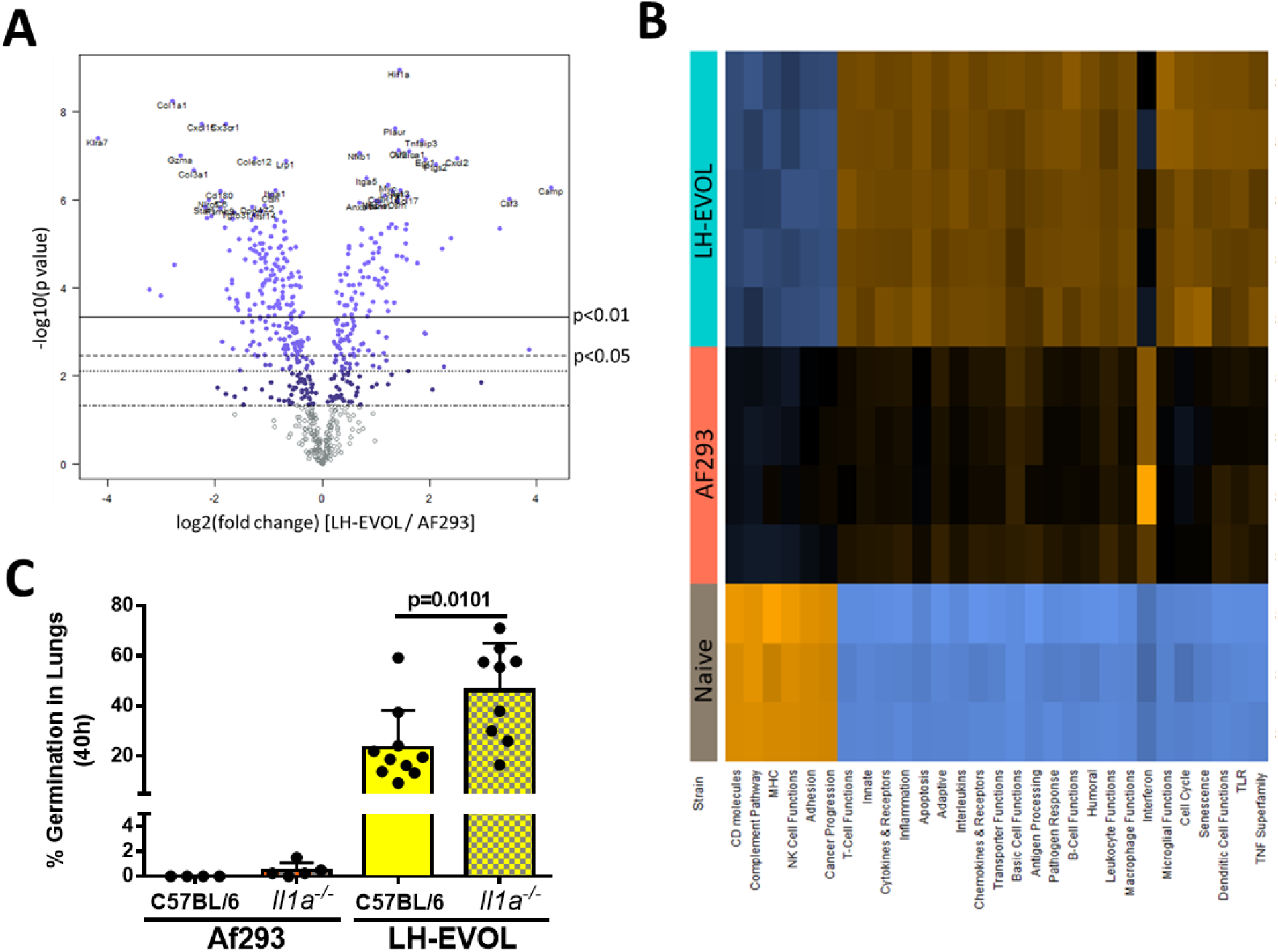
Differential pulmonary inflammatory gene expression and immune response after AF293 and LH-EVOL challenge. C57BL/6 mice were inoculated with either PBS or 4×10^7^ AF293 or LH-EVOL conidia and euthanized at 40 h post-inoculation. Total mRNA was extracted from the whole lung, and gene expression was analyzed by NanoString nCounter PanCancer Immune Profiling panel. (A) Volcano plot showing the distribution of fold changes in gene expression in LH-EVOL challenged mice compared to AF293 challenged mice. (B) Pathway signature of mice challenged with either PBS, AF293, or LH-EVOL. Orange color indicates a pathway is enriched, while blue color indicates a pathway is decreased. (C) C57BL/6 and IL-1α mice were inoculated with either 4×10^7^ conidia of either AF293 or LH-EVOL. At 40 h post-challenge mice were euthanized and lungs saved for histological analysis. Formalin-fixed lungs were paraffin embedded, sectioned, and stained with GMS for microscopic analysis of fungal germination. Data are pooled from two independent experiments. Statistical significance was determined using a Mann–Whitney U test.

**Table 1.** Select gene transcripts upregulated following AF293 challenge.

| **GENE**<br>**NAME** | **FOLD CHANGE**<br>**(LOG2)** | **BY P-VALUE** |
| --- | --- | --- |
| *Cxcl9* | 2.92 | 0.0155 |
| *Cxcl10* | 2.68 | 0.000138 |
| *Gzma* | 2.57 | 0.000536 |
| *Stat1* | 2.11 | 0.000174 |
| *Angpt1* | 2.07 | 0.00212 |
| *Nlrc5* | 2.04 | 0.00163 |
| *Zbp1* | 1.73 | 0.00243 |
| *Ddx60* | 1.66 | 0.00553 |
| *Gbp5* | 1.52 | 0.0273 |
| *Irf7* | 1.35 | 0.00715 |
| *Isg15* | 1.32 | 0.0182 |
| *Ifit3* | 1.27 | 0.0357 |
| *Ddx58* | 0.664 | 0.0166 |
| *Ifih1* | 0.296 | n.s. |

**Table 2.** Select gene transcripts upregulated following LH-EVOL challenge.

| **GENE**<br>**NAME** | **FOLD CHANGE**<br>**(LOG2)** | **BY P-VALUE** |
| --- | --- | --- |
| *Camp* | 4.34 | 0.000918 |
| *Csf3* | 3.57 | 0.000115 |
| *Cxcl2* | 2.59 | 0.000423 |
| *Cxcl1* | 2.47 | 0.000918 |
| *Il6* | 2.32 | 0.00212 |
| *Ptgs2* | 2.20 | 0.000369 |
| *Hif1a* | 1.51 | 0.000249 |
| *Csf2* | 1.51 | 0.000369 |
| *Cxcl5* | 1.42 | 0.00515 |
| *Il12a* | 1.23 | 0.00442 |
| *Nfkbia* | 1.08 | 0.000139 |
| *Il1a* | 1.06 | 0.000369 |
| *Nfkb1* | 0.77 | 0.000369 |
| *Il1b* | 0.66 | 0.00139 |

The marked elevation of *Il1a* transcripts observed following LH-EVOL challenge fits with our previous observation that IL-1α is essential for host resistance against highly virulent isolates of *A. fumigatus* that can rapidly germinate in the lung airways^4,42^. To test the importance of IL-1α in host resistance against the LH-EVOL strain, C57BL/6J mice and *Il1a*^(−/−)^ mice were challenged with either AF293 or LH-EVOL. At 40 hours post-challenge, lungs were collected for histological analysis to determine fungal germination. We found that not only did C57BL/6J and IL-1α-deficient mice challenged with LH-EVOL have significantly greater fungal germination than AF293 challenged mice (Figure 2C), but IL-1α-deficient mice challenged with LH-EVOL also had significantly more fungal germination than C57BL/6J mice challenged with LH-EVOL (Figure 2C). Taken together these data reveal that the rapid germinating LH-EVOL strain is markedly more inflammatory.

### Genomic analysis of LH-EVOL identifies a potential truncation mutation within *Afu5g08390* (*sskA*)

Selective serial culturing of an organism will often result in genetic modifications as the organism responds to a given environmental pressure, such as growth in the lung microenvironment without direct *a priori* knowledge about the exact nutrients that are biologically available for fungal growth. Therefore, we asked what genomic changes arose in LH-EVOL to determine the genetic basis for its rapid germination phenotype. Three single spore clones of LH-EVOL were selected for further analysis. Each of the clones of LH-EVOL were able to germinate substantially faster than the parental AF293 strain in LH medium in the *in vitro* germination assay (Figure 3A). Next, whole genome sequencing of these three clones of LH-EVOL was conducted and compared with its parental AF293 strain. Whole genome sequencing revealed conserved INDEL mutations in *Afu5g08390* across the three LH-EVOL clones (Figure 3B). *Afu5g08390* encodes for SskA which is a response regulator in the high osmolarity glycerol (HOG) pathway^43^. These INDEL mutations are predicted to insert premature stop codons in the *Afu5g08390* gene that may induce premature truncation before the recognition (REC) domain (Figure 3C), a domain necessary for the signaling function of Ssk1, the SskA homologue, in yeast^29^.

**Figure 3.**
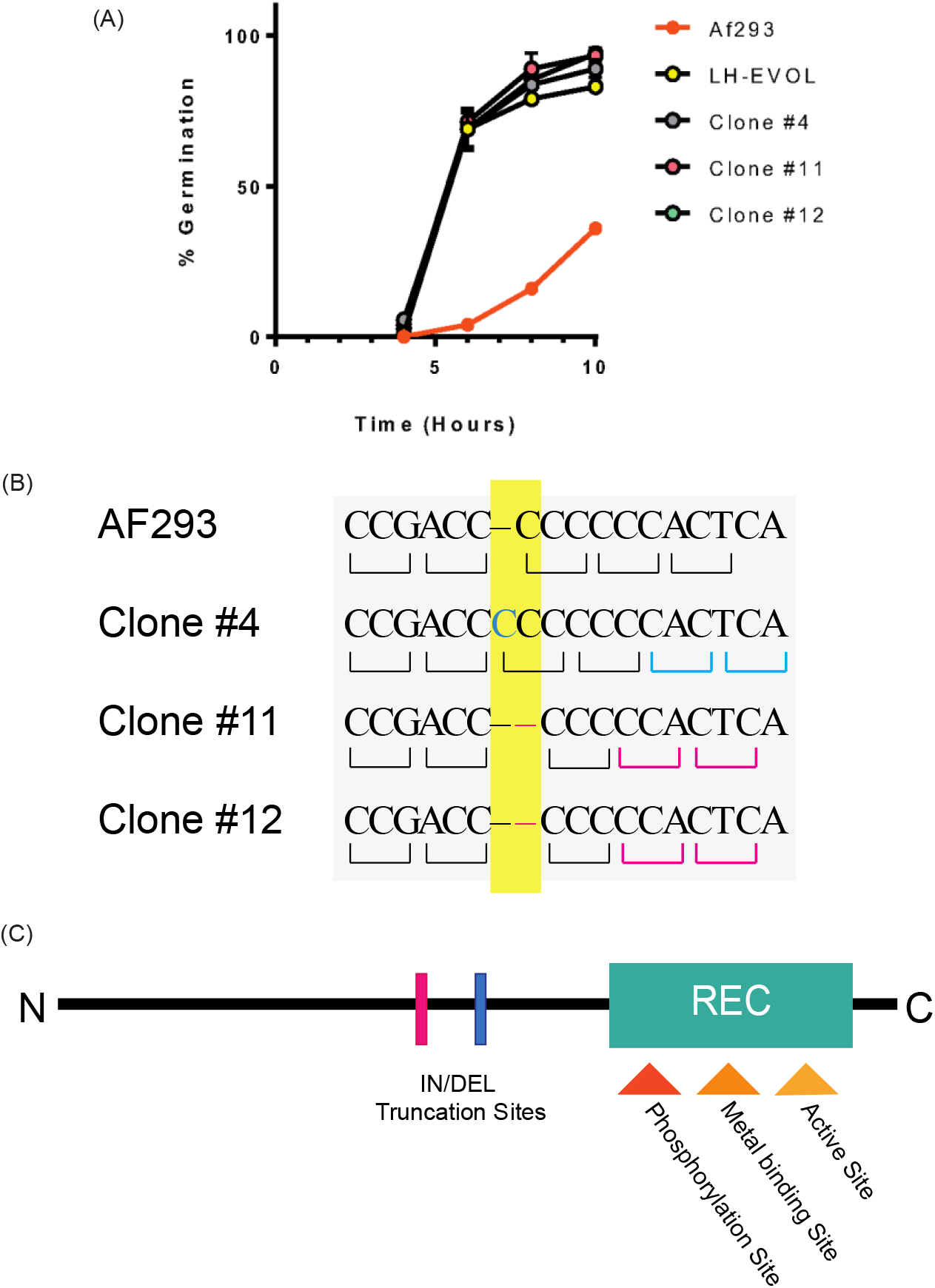
Identification of INDEL mutations in the *Afu5g08390* (*sskA*) gene of *A. fumigatus*. (A) Percent germination over time in lung homogenate media of AF293, the bulk LH-EVOL strain, or individual LH-EVOL clones (clone #4, clone #11, or clone #11). Data are representative of at least 2 independent experiments consisting of 3 biological replicates per group. Each symbol represents the group mean ± 1 standard error of the mean. Statistical significance was determined using one-way ANOVA with Dunn’s test. (B) DNA-sequencing of LH-EVOL clones #4, #11, and #12 revealed INDEL mutations in the *Afu5g08390* (*sskA*) gene. (C) Based on the predicted amino acid changes due to the frameshift caused by these INDEL mutations it is predicted that a premature stop codon causes the truncation of SskA prior to its REC domain in the LH-EVOL strains.

### LH-EVOL has a decreased SakA-dependent response under osmotic stress

SskA is a response regulator within the high osmolarity glycerol pathway which drives the activation of the MAPKKK SskB, MAPKK PbsB, and MAPK SakA^44,45^. SakA is critical for maintaining homeostasis in *A. fumigatus* during osmotic stress^13,46,47^. Based on the predicted truncation of SskA prior to its REC domain (Figure 3C), we hypothesized that the LH-EVOL strain has a decreased SakA signaling response. To directly test this AF293, AF293 Δ*sakA*, and LH-EVOL were inoculated onto GMM plates and GMM plates supplemented with 1M NaCl, 1M KCl, 0.5M CaCl_2_, and 1M Sorbitol for 72 hours at 37°C (Figure 4A). LH-EVOL grown on GMM formed significantly larger colonies than AF293, similar to AF293 Δ*sakA* (Figure 4B). Importantly, when LH-EVOL was grown on GMM plates supplemented with osmotic stressors, its growth was substantially altered compared with AF293, similar to the response of the AF293 Δ*sakA* mutant (Figure 4C)^13^. Specifically, both LH-EVOL and AF293 Δ*sakA* were significantly impaired in their ability to grow in the presence of 1M NaCl, while growing significantly better in the presence of 1M sorbitol.

**Figure 4.**
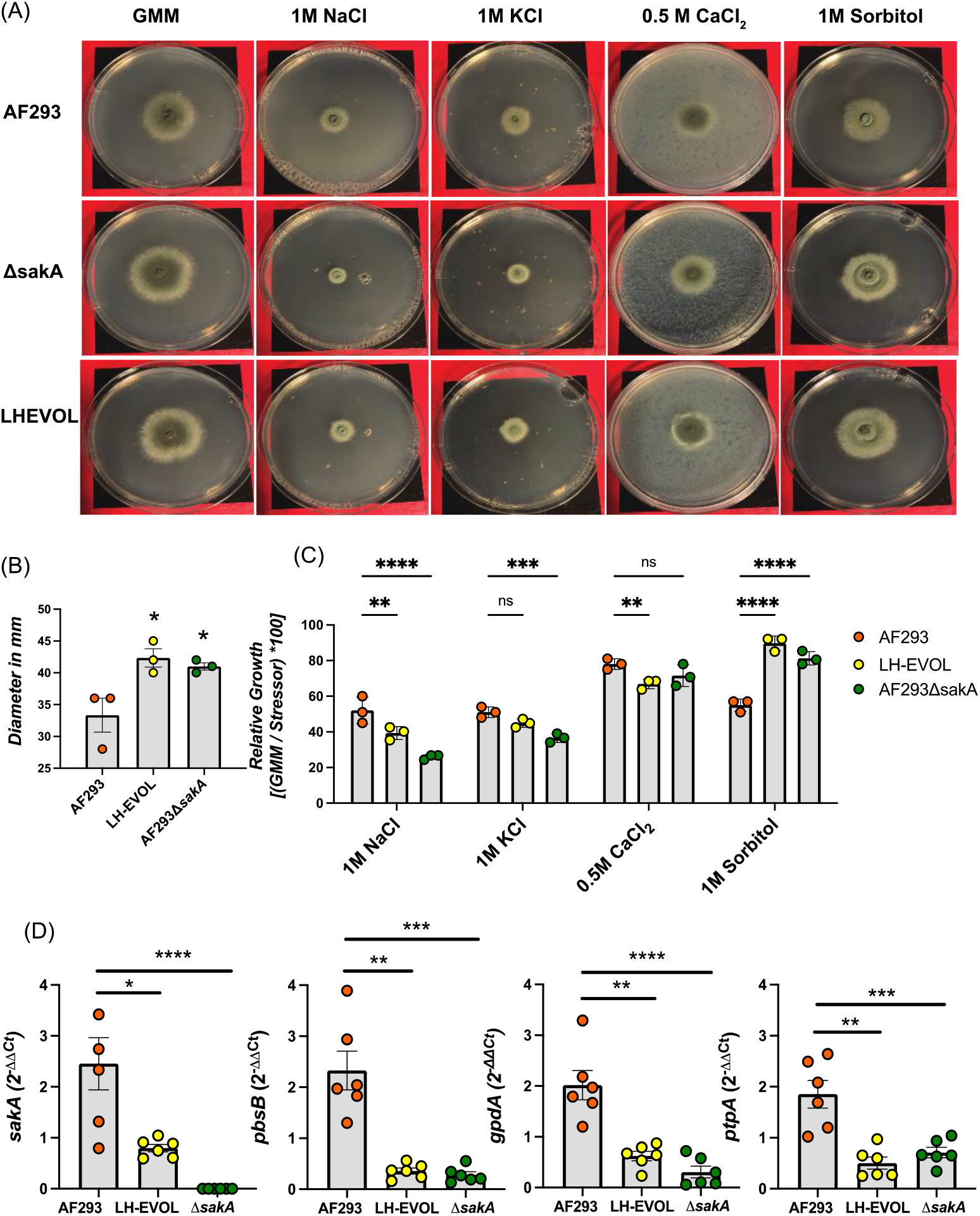
LH-EVOL strain demonstrates deficiencies in *sakA*-dependent osmotic stress response. (A) AF293, AF293 Δ*sakA*, and LH-EVOL conidia were inoculated on GMM plates or GMM plates supplemented with 1M NaCl, 1M KCl, 0.5M CaCl_2_, or 1M Sorbitol and incubated for 72 hours at 37°C. (B) Diameter of colonies grown on GMM for 72 hours in millimeters (mm). Diameter of the colony was used to calculate the percent of inhibited growth compared to its respective measurement on GMM. (D) A 30-hour biofilm was transferred to fresh liquid GMM media supplemented with 1M NaCl for 30 minutes at 37°C. Quantitative RT-PCR was used to measure the change in transcript production of *sakA* and *sakA*-dependent genes *pbsB, gpdA*, and (E) *ptpA*. Ct values were initially adjusted to *actA* for ΔCt, then 1M NaCl was compared to GMM for ΔΔCt values. Data are pooled of results from 2 independent experiments. Statistical significance was assessed using (B) a one-way ANOVA with Tukey’s test, (C) a two-way ANOVA and (D) one-way ANOVA with a Kruskal-Wallis test. (*) indicates a p-value ≤ 0.05; (**) indicates a p-value ≤ 0.01; (***) indicates a p-value ≤ 0.001.

Since LH-EVOL appears to have an altered response to osmotic stress similar to AF293 Δ*sakA*, we next explored the transcriptional responses of AF293, AF293 Δ*sakA*, and LH-EVOL conidia to 1M NaCl, which is known to drive SakA activation and signaling^13^. Each strain was grown in GMM media for 30 hours at 37°C before being transferred to either GMM or GMM supplemented with 1M NaCl for 30 minutes. We choose to examine *sakA* expression itself, as well as SakA-dependent transcripts *ptpA*, *gpdA*, and *pbsB* which are highly upregulated in AF293 but not in AF293 Δ*sakA* following osmotic stress with 1M NaCl^13,48^. Transcript levels of *sakA, ptpA, gpdA*, and *pbsB* were all increased to a significantly greater extent in parental strain AF293 as opposed to LH-EVOL and AF293 Δ*sakA* when exposed to 1M NaCl (Figure 4D). These data support the hypothesis that the allele of *Afu5g08390* (*sskA*) found in the LH-EVOL strain likely encodes a SskA truncation mutant with decrease signal transduction.

### Loss of *Afu5g08390* (*sskA*) in AF293 results in increased germination

Since the INDEL mutations found within *Afu5g08390* (*sskA*) in the LH-EVOL strain are predicted to result in a premature truncation mutation (Figure 3C), which resulted in decreased SakA-dependent transcriptional response to osmotic stress (Figure 4), we next tested whether the complete loss of SskA results in increased germination potential in AF293. To do this, we constructed a *sskA* null mutant using homologous recombination in the AF293 strain of *A. fumigatus*, as well as an ectopic complementation strain (Supplemental Figure 1). When cultured on GMM plates the AF293 Δ*sskA* strain had significantly greater radial growth than the parental AF293, which was similar to what was observed with the AF293 Δ*sakA* and LH-EVOL strains (Supplemental Figure 2A). Next, to determine if the AF293 Δ*sskA* strain had decreased SakA signaling we explored the transcriptional responses of AF293, AF293 Δ*sakA*, and AF293 Δ*sskA* biofilms to 1M NaCl. Each strain was cultured in GMM media for 30 hours at 37°C before being transferred to either GMM or GMM supplemented with 1M NaCl for 30 minutes. Similar to previous results (Figure 4D), transcript levels of *sakA, ptpA, gpdA*, and *pbsB* were all increased in AF293 relative to AF293 Δ*sakA* when exposed to 1M NaCl (Supplemental Figure 2B-E). Importantly, AF293 Δ*sskA* failed to show elevated transcript levels of *sakA, ptpA, gpdA*, and *pbsB*, which supports the hypothesis that SskA-SakA signaling is decreased in the AF293 Δ*sskA* strains (Supplemental Figure 2B-E).

Next, to determine if AF293 Δ*sskA* was able to germinate faster than AF293 or the complemented AF293 Δ*sskA^RC^* strain, we conducted an *in vitro* germination assay in lung homogenate medium. When AF293 Δ*sskA* was grown in lung homogenate media, robust germination was observed compared to the parental strain or the complemented AF293 Δ*sskA^RC^* (Figure 5A). In contrast, all three strains germinated at equivalent rates in nutrient rich GMM + YE medium (Figure 5B), demonstrating a lack of intrinsic changes to the overall germination potential of the AF293 Δ*sskA* strain. To determine if AF293 Δ*sskA* also has an increased germination rate during initial airway infection of mice, C57BL/6J mice were inoculated intratracheally with either AF293, AF293 Δ*sskA*, and AF293 Δ*sskA^RC^*. Twelve hours post inoculation, AF293 Δ*sskA* had germinated to a significantly greater degree than either the AF293 or AF293 Δ*sskA^RC^* (Figure 5C). Thus, loss of SskA signaling in AF293 strain of *A. fumigatus* results in enhanced germination potential within the lung microenvironment.

**Figure 5.**
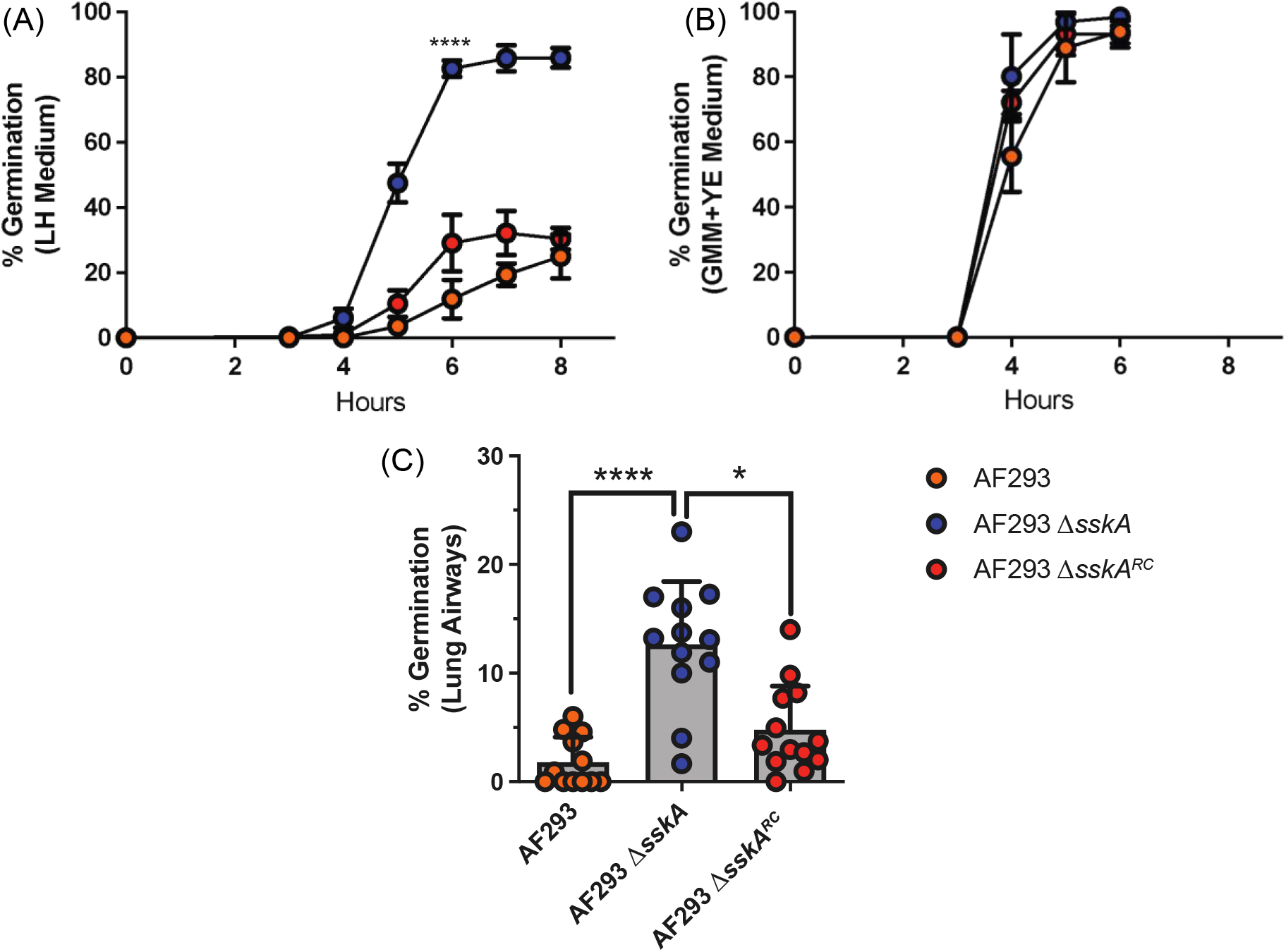
Loss of *sskA* in AF293 increases germination rate. Conidia germination of AF293, AF293 Δ*sskA*, and AF293 Δ*sskA^RC^* was determined by wet mount by microscopically counting the numbers of conidia and germlings in either lung homogenate media (A) or 1% GMM supplemented with yeast extract (B). Data are representative of at least 2 independent experiments consisting of 3 biological replicates per group, Each symbol represents the group mean ± 1 standard error of the mean. (C) *In vivo* conidia germination in airways of C57BL/6J mice with 4×10^7^ conidia of AF293, AF293 Δ*sskA*, and AF293 Δ*sskA^RC^*. At 12 hpi, mice were euthanized, and BALF was collected to quantify germination in the airways. Data are pooled of results from 2 independent experiments. Statistical significance was assessed using an (A) unpaired t-test or (C) one-way ANOVA with a Dunn post-test. (*) indicates a p-value ≤ 0.05; (***) indicates a p-value ≤ 0.001.

### Loss of the MAPKs *SakA* and *MpkC* results in increased germination

SakA associates with MpkC, a MAPK of the cell wall integrity pathway, in the nucleus^46,49^. Therefore, we wanted to test the role of both the SakA and MpkC MAPK proteins in the regulation of germination of AF293 in the pulmonary environment. To determine if SakA and MpkC regulate *A. fumigatus* germination, the rate of conidial germination of AF293 Δ*sakA* and AF293 Δ*mpkC* mutant strains were assessed in lung homogenate medium. We found that germination indeed increased in both AF293 Δ*sakA* and AF293 Δ*mpkC* relative to the parental strain (Figure 6A). Additionally, the germination rate of both AF293 Δ*sakA* and AF293 Δ*mpkC* were similar to what we had observed for LH-EVOL and AF293 Δ*sskA* (Figure 1B and Figure 4A, respectively). In contrast, all strains have similar germination rates in GMM + YE medium, demonstrating that there were no global increases to the overall germination potential of these mutants (Figure 6B). To determine if AF293 Δ*sakA* and AF293 Δ*mpkC* also have an increased germination rate during initial airway infection of mice, C57BL/6J mice were inoculated intratracheally with either AF293, AF293 Δ*sakA*, or AF293 Δ*mpkC*. Twelve hours post inoculation, both AF293 Δ*sakA* and AF293 Δ*mpkC* had germinated to a significantly greater degree than their parental strain (Figure 6C). Thus, SskA signaling appears to converge on both the SakA and MpkC MAPKs in the AF293 strain of *A. fumigatus* to regulate germination within the lung microenvironment.

**Figure 6.**
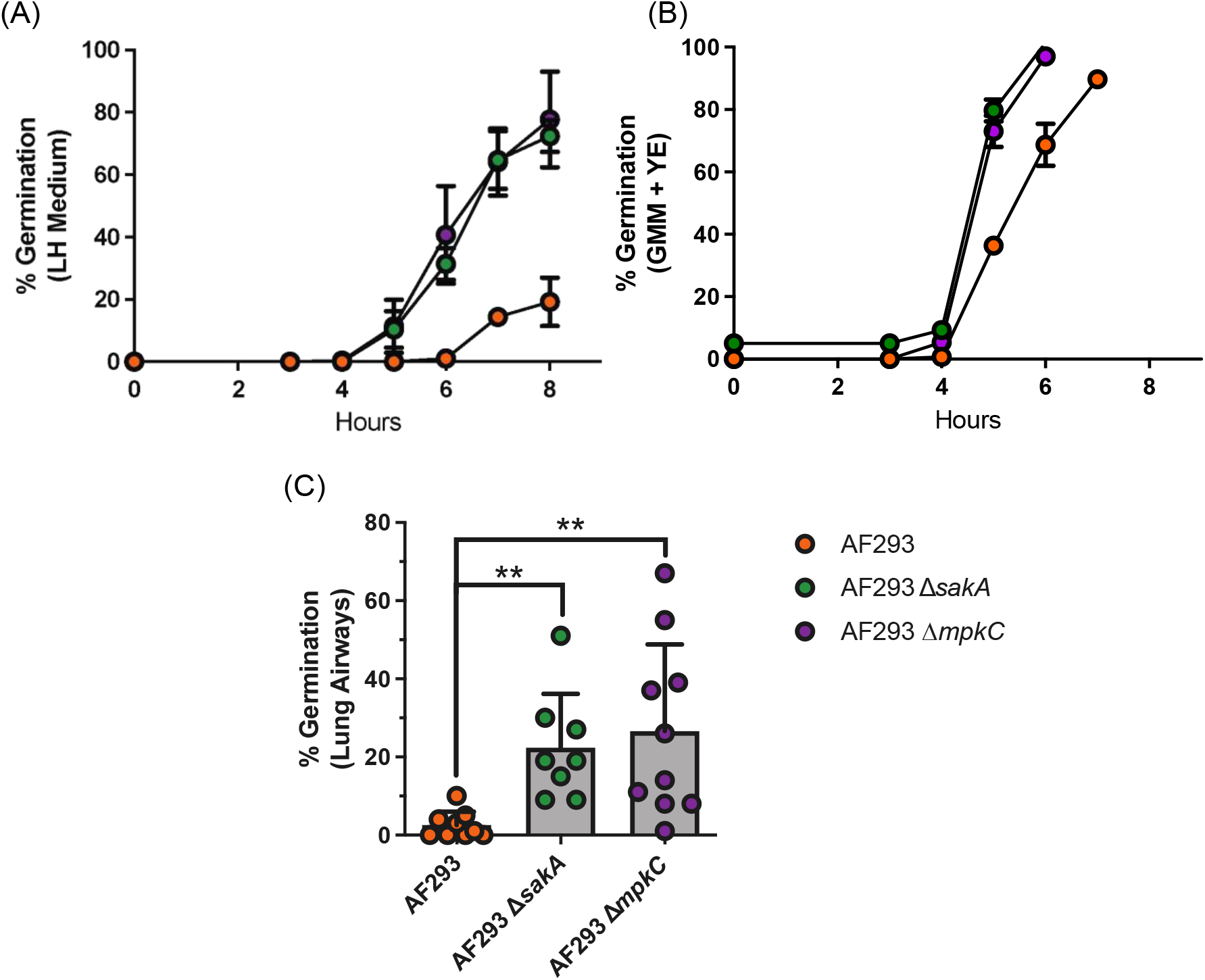
SakA- and MpkC-null mutants have increased conidia germination kinetics *in vitro* and *in vivo*. (A-B) Conidia germination of AF293, AF293 Δ*sakA*, and AF293 Δ*mpkC* was determined by wet mount by microscopically counting the numbers of conidia and germlings in either lung homogenate media (A) or 1% GMM supplemented with yeast extract (B). Data are representative of at least 2 independent experiments consisting of 3 biological replicates per group. Each symbol represents the group mean ± 1 standard error of the mean. (C) *In vivo* conidia germination in airways of C57BL/6J mice with 4×10^7^ conidia of AF293, AF293 Δ*sakA*, and AF293 Δ*mpkC*. At 12 hpi, mice were euthanized, and BALF was collected to quantify germination in the airways. Data are pooled of results from 2 independent experiments. Statistical significance was assessed using an (A) unpaired t-test or (C) one-way ANOVA with a Dunn post-test. (**) indicates a p-value ≤ 0.01.

### Protein sequence analysis of the SakA signaling components in CEA10 identifies differences in TcsB and MpkC

As the CEA10 strain of *A. fumigatus* is able to rapidly germinate both *in vitro* in lung homogenate medium and *in vivo* in murine lungs^4^, we explored CEA10’s transcriptional response to 1M NaCl in the biofilm transfer assay. Specifically, CEA10 was cultured in GMM for 30 hours at 37°C before being transferred to either GMM or GMM supplemented with 1M NaCl for 30 minutes. While AF293 had elevated transcript levels of *sakA, ptpA, gpdA*, and *pbsB*, CEA10 did not have increased levels of these transcript (Supplemental Figure 2). These data support the hypothesis that SskA-SakA signaling is altered in CEA10 relative to AF293.

Next, we compared the SskA amino acid sequences between AF293 and CEA10. Using BLASTp we found that the SskA protein sequences where 100% identical between AF293 and CEA10 (Supplemental Table 4). Next, we expanded our BLASTp analysis to include all the known components of the SskA-SakA signaling pathway. This analysis revealed that YpdA, SskB, PbsB, and SakA were also 100% identical between Af293 and CEA10 (Supplemental Figure 2). In contrast, BLASTp sequence alignment of the putative histidine kinase response regulator TcsB, homologue to *S. cerevisiae* SLN1 and integral to the activation of Ssk1 in yeast^44,50,51^, revealed a 13 amino acid deletion (AA 709-721, XP_001481640.1) within the predicted ATPase (AA 696-863, XP_001481640.1) domain in the histidine kinase domain of the CEA10 TcsB protein that is not present in AF293 (Supplemental Figure 2). Additionally, BLASTp sequence alignment of MpkC, which can interact with SakA signaling^46,49,52,53^, shows 4 non-synonymous mutations (Supplemental Table 4). Thus, our BLASTp analysis suggest that changes in the SskA protein of CEA10 are likely not responsible for the increased germination rate of that strain, while change in other proteins in the SakA signaling network, particularly TcsB, might be functionally different.

### SskA-SakA signaling limits *A. fumigatus* germination in decreased glucose conditions

The ASL which lines the airway epithelium including the lungs is a key component of respiratory homeostasis and facilitates mucociliary clearance of foreign particulates and pathogens. The glucose concentration of ASL is approximately 0.4 m*M*, substantially lower than typical *in vitro* culturing conditions. In yeast, glucose starvation can mediate activation of the stress MAPK Hog1 in a Ssk1-dependent manner^53^. Consequently, we hypothesized that limited glucose availability may limit AF293 germination through SskA-SakA activation. To test this hypothesis, an *in vitro* germination assay using the AF293, LH-EVOL, AF293 Δ*sskA*, AF293 Δ*sakA*, and AF293 Δ*mpkC*, strains was conducted in either 1% GMM (55.56 mM of glucose) or 0.25% GMM (13.89 mM of glucose). Interestingly, we found that while AF293 underwent limited germination in 1% GMM it nearly completely failed to germinate in 0.25% GMM (Figure 7A); however, the AF293 Δ*sskA* (Figure 7C), AF293 Δ*sakA* (Figure 7D), and AF293 Δ*mpkC* (Figure 7E) underwent greater germination in 0.25% GMM than in 1% GMM. When we examined LH-EVOL we found that the germination rate in both 1% GMM and 0.25% GMM were equivalent (Figure 5B). Thus, it appears that in the absence of SskA-SakA signaling, there is an increased potential for *A. fumigatus* strain to germinate under low glucose conditions which would be found in the airways upon initial deposition.

**Figure 7.**
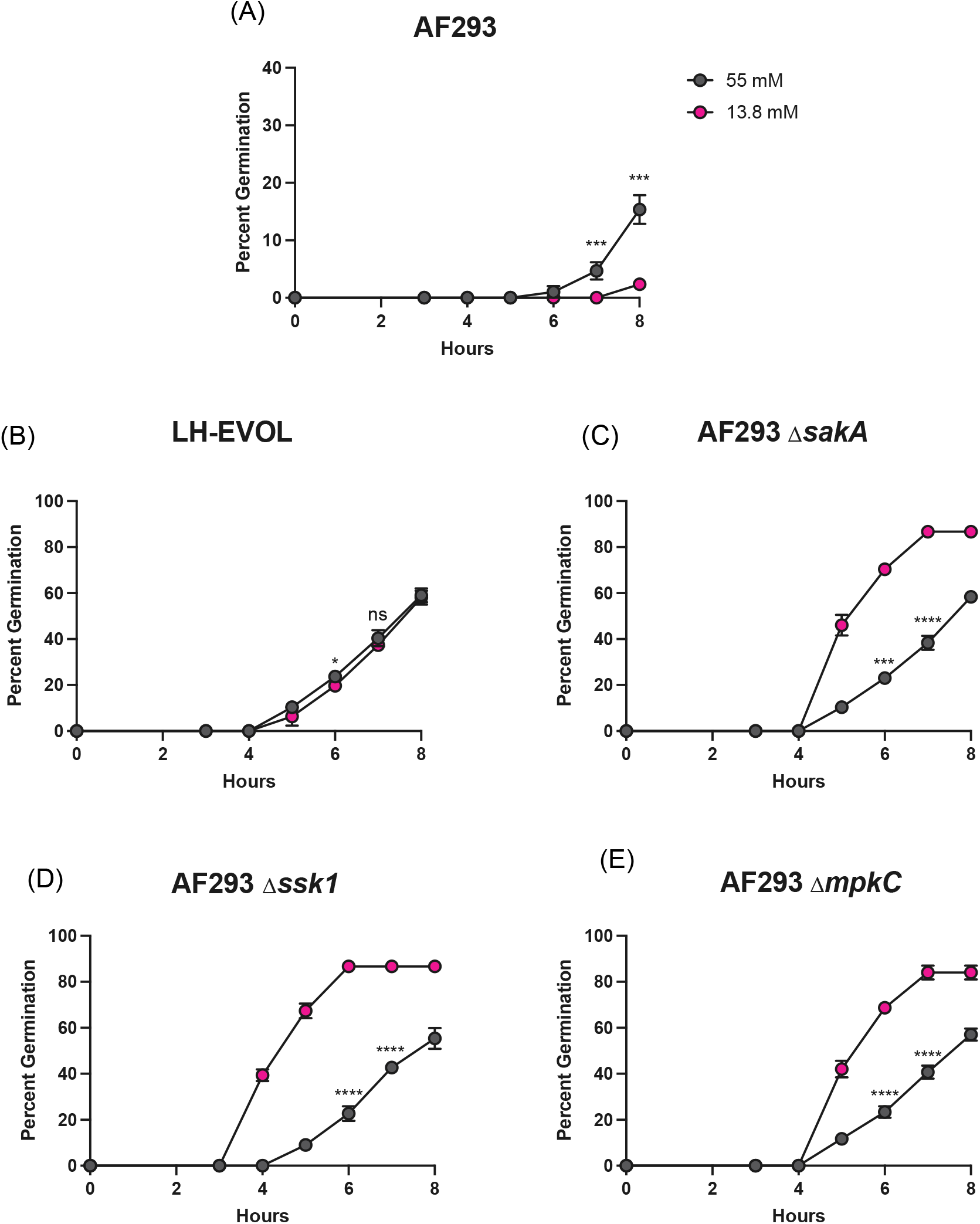
Increased germination rate of lung-adapted *A. fumigatus* strains in low glucose conditions. Conidia germination of AF293 (A), LH-EVOL (B), AF293 Δ*sakA* (C), AF293 Δ*sskA*, and AF293 Δ*mpkC* (E) was determined by wet mount and microscopically counting the numbers of conidia and germlings in either 1% GMM or 0.25% GMM medium. Statistical significance was assessed using an unpaired Student’s t-test. (*) indicates a p-value ≤ 0.05; (**) indicates a p-value ≤ 0.01; (***) indicates a p-value ≤ 0.001; (****) indicates a p-value ≤ 0.0001.

## DISCUSSION

After entering the airways of a mammalian host, conidia are typically cleared by mucociliary clearance or innate phagocytotic responses^1–3,54^. In order to drive disease within a host, two types of virulence traits have been proposed: (1) disease initiating factors, which are factors that either promote the initiation of infection or disease pathology; and (2) disease progressing factors, which facilitate microbe persistence, host damage, and disease progression in an established infection microenvironment^55^. Our previous work has demonstrated early fungal germination within the airway space is a disease initiating trait in a model of *A. fumigatus* induced bronchopneumonia^4^. However, the molecular mechanism(s) responsible for driving the increased fungal germination rate of certain *A. fumigatus* isolates remains elusive. In this study, we took an experimental serial passage approach to identify factors that may drive enhanced fungal germination. Through this approach we isolated a novel, lung-adapted strain, LH-EVOL, from the parental AF293 strain. The LH-EVOL strain showed increased germination *in vivo* in murine airways and *in vitro* in lung homogenate medium (Figure 1) in addition to enhanced proinflammatory cytokine responses (Figure 2).

Whole genome sequencing of LH-EVOL revealed the presence of INDEL mutations in SskA that are predicted to result in the truncation of the SskA recognition (REC) domain^37^ (Figure 3), a domain necessary for SskA to dimerize with SskB to phosphorylate PbsB. In support of a predicted truncation of SskA’s REC domain, the LH-EVOL strain does not induce as strong of a SakA-dependent response as measured by transcript abundance of SakA target genes during osmotic stress (Figure 4); thus, the *sskA* allele identified in the LH-EVOL strain likely encodes an altered or non-functional SskA protein. However, one limitation of this study is that the function of the SskA protein found in LH-EVOL has not been directly determined. The prediction that the LH-EVOL allele of *sskA* result in a non-functional protein is, however, further supported by the finding that AF293 Δ*sskA* also germinates more rapidly than the parental strain *in vitro* in lung homogenate medium (Figure 5A) and *in vivo* in the murine airways (Figure 5C) which is what was observed in the LH-EVOL strain (Figure 1). SskA is part of the SakA MAPK stress response pathway and has been shown to be important for SakA activation in the Afs35 strain of *A. fumigatus*^26^. During germination of *A. nidulans*, SakA is dephosphorylated and exits the nucleus^49^. Based on the location of the predicted premature stop codon due to an INDEL mutation found in the *sskA* gene of LH-EVOL (Figure 3), it can be predicted that there would be less phosphorylated SakA in LH-EVOL and AF293 Δ*sskA*, making those strains poised to germinate quickly within the lung airways. This is further supported by our data demonstrating that AF293 Δ*sakA* also exhibits increased germination rates *in vitro* in lung homogenate medium (Figure 6A) and *in vivo* in the airways (Figure 6C). Interestingly, AF293 Δ*sakA* appears to undergo greater germination potential in the murine lungs than AF293 Δ*sskA*, although this was not compared directly. In *S. cerevisiae* Hog1 signaling has two branches for its activation: Sln1 and Sho1 which are Ssk1-dependent and Ssk1-independent, respectively^56^, which also seems to be case in *A. fumigatus*^57^. Our study only examined the role of the SskA-dependent branch in *A. fumigatus* germination, but future studies should examine the interplay of these two SakA signaling branches in regulating *A. fumigatus* germination.

As CEA10 is also a strain of *A. fumigatus* that can rapidly germinate in the lungs^4^, we next examined the SskA-SakA signaling network in that strain. Interestingly, compared with AF293, CEA10 had a significantly weaker induction of SakA-dependent transcripts (Supplemental Figure 2). However, the amino acid sequence identity of SskA and SakA where 100% between AF293 and CEA10. Upstream of SskA the histidine kinase TcsB, a homologue of SLN1 in yeast that is necessary for Ssk1 and HOG1 activation^44,50,51^, in CEA10 has a 13 amino acid deletion in the predicted ATPase domain (Supplemental Figure 3). This deletion could account for reduced expression of *pbsB*, *sakA* and SakA dependent genes in CEA10 during osmotic stress (Supplemental Figure 2), but this needs to be experimentally further explored. Therefore, strains of *A. fumigatus* with decreased activity of SakA appear to have an increased potential for rapid germination and growth in murine lungs. Future studies should look at the relationship of *A. fumigatus* germination and the activity of specific SakA-dependent genes.

Within the pulmonary environment, the specific nutrient(s) driving *A. fumigatus* germination remains elusive. The airways of the lungs are a low-nutrient environment. Recent *in vivo* transcriptional analysis of the response of *A. fumigatus* in the murine airways demonstrates that *A. fumigatus* undergoes significant stress from iron, zinc, and nitrogen starvation^58^. In our experiments both the LH-EVOL strain and AF293-based SskA null mutant have an increased ability to germinate in 1% GMM when compared to their parental AF293 strain (Figure 5). Furthermore, the AF293-based SakA-null mutant has an increased ability to germinate in 1% GMM^13^. Interestingly, the ability of wild-type AF293 to germinate can be rescued by the supplementation of GMM with yeast extract (Figure 5B) or complete medium containing yeast extract, peptone, and tryptone^13^. This is significant because it has previously been shown that AF293 conidia are unable to utilize nitrate, the nitrogen source available in GMM media, as effectively as AF293 Δ*sakA* for growth in a SakA-dependent manner^13^. Taken together these data suggest that *A. fumigatus* may need to be able to effectively utilize non-preferred nitrogen sources within the lung microenvironment to initiate germination, which will be explored in future studies.

Glucose levels in ASL of a healthy adult are approximately 0.4 m*M* ^14^. These levels are ~10-fold reduced when compared with plasma^14^ and is thought to limit microbial growth thus limiting infections^15^. Surprisingly, we found that the LH-EVOL and AF293-based SskA, SakA, and MpkC null mutants were still able to germinate in 0.25% GMM, whereas the parental AF293 isolate was not. SakA has previously been shown to be directly involved in glucose uptake, glycogen and trehalose storage, and trehalose utilization by *A. fumigatus*^59^. In addition, SakA-dependent glucose sensing modulates biofilm development^60^. Furthermore, in *Cryptococcus neoformans*, Hog1 and protein kinase A (PKA) have been shown to be critical for adaptation to the glucose-limited environment of the lungs and glucose-replete environment of the brain, respectively^61^. In low glucose conditions, Hog1 of *C. neoformans* becomes activated, resulting in translational repression through decreased ribosome biogenesis which is necessary for stress adaptation to the low glucose environment of the lungs. In *A. fumigatus*, PKA signaling, regulated by SakA and MpkC^59^, has been shown to be critical for growth in low glucose environments and conidial germination^62^. Considering these findings, our data would predict that the low glucose environment of the lungs might similarly drive SakA activation in some *A. fumigatus* strains, such as AF293, limiting germination but increasing resistance against physiological stressors. In contrast, rapidly germinating strains may be more susceptible to osmotic and oxidative stressors in the host environment due to decreased SakA activity. This fits with the observation that CEA10 is more rapidly cleared than AF293 in a zebrafish model of infection^5^. Consequently, SakA could be particularly important for *A. fumigatus* to adapt to the cystic fibrosis (CF) lung environment where there is increased oxidative stress^63^ and ASL glucose concentrations can be 10 times higher than healthy adults^16^. Ross *et al* (2021) demonstrated a distinct relationship between activation of the SakA pathway and adaptation by *A. fumigatus* to the lung environment in a CF patient^48^. In that study, PbsB, an intermediary kinase between SskA and SakA, was found to have a missense mutation that resulted in increased SakA activity in response to osmotic stress^48^. These data suggest a potentially important and distinct relationship between SakA activity in *A. fumigatus* in response to nutritional and osmotic stressors in the pulmonary environment, which may be critical to understanding disease initiation and progression within both acute and chronic models of aspergillosis.

Finally, our inflammatory profiling of pulmonary responses to both AF293 and LH-EVOL reinforces our knowledge that early germination in lung airways is a key driver of the distinct immunopathological phenotypes observed within different *A. fumigatus* isolates, where fast germinating isolates require an IL-1α dependent host response to maintain host resistance^4^ (Figure 2C). Our inflammatory profiling also found that *Hif1a* mRNA was expressed to a greater extent following LH-EVOL challenge than AF293 (Table 2), fitting with our previous observation that HIF1α-dependent inflammatory responses are necessary for host resistance against the rapidly germinating CEA10 strain of *A. fumigatus*, but not AF293^40^.

Notably, we found a stronger IFN signaling signature following challenge with AF293 than LH-EVOL (Figure 2B). Recent work has demonstrated a critical role for both type I and type III IFNs in driving host resistance against *A. fumigatus*^64^. Based on our NanoString analysis (Figure 2), slow germinating strains of *A. fumigatus* may be more potent inducers of the type I and/or type III IFN response, which warrants further exploration. In support of this, human bronchial epithelial cells respond to resting conidia, rather than swollen conidia or hyphae, by enhanced secretion of IFN-β and CXCL10^41^. The expression of type I IFN (IFN-α and IFN-β) and CXCL10 can be induced by *A. fumigatus* dsRNA through both TLR3-Trif and MDA5-MAVS dependent signaling^41,65,66^. Overall, this suggests that the induction of IFN may occur in response to early development stages of *A. fumigatus* infection, while the robust pro-inflammatory response (e.g. IL-1α release) is driven by the later invasive hyphal stage. Differential host response to other fungal infections over time have been previously described. Recent work examining the inflammatory response of vaginal epithelial cells to *Candida* spp. demonstrate an early, homogeneous type I IFN response to all *Candida* spp., but at later time points, the inflammatory response diverges in a species specific and damage-dependent manner^67^.

In conclusion, this study further emphasizes the strain-specific virulence, pathology, and inflammatory responses that occur during *A. fumigatus* infections. Our study also identifies a role for the loss of SakA MAPK signaling in enhancing conidial germination, particularly in low glucose environments. This likely comes with the recompense of decreased resistance to osmotic and oxidative stress, which could be critical to disease progression during chronic fungal diseases^67^. Thus, future studies must address the role of these traits and pathways in both acute and chronic models of aspergillosis.

## AUTHORS CONTRIBUTIONS

Conceived the experiments: MEK, RAC, JJO. Designed the experiments: MEK, RMT, AKC, CHK, BSR, RAC, JJO. Performed the experiments: MEK, MS, RMT, AKC, BSR. Generated fungal mutants: CHK, DM. Analyzed the data: MEK, MS, AKC, CHK, DM, LAL, JES, RAC, JJO. Wrote the paper: MEK, JJO.

## ACKNOWLEDGEMENTS

Thank you to Drs. Deborah Hogan and James Bliska (Geisel School of Medicine at Dartmouth) for helpful discussion on this project and manuscript. The authors thank Dr. Charles Puerner (Dartmouth College, Cramer Lab) for quantifying the glucose levels in our lung homogenate medium. The authors thank Dr. Jong Heon Kim (USDA ARS, Albany, CA) for providing the AF293 Δ*sakA* and AF293 Δ*mpkC* strains. Research in this study was supported in part by institutional startup funds to JJO in part through the Dartmouth Lung Biology Center for Molecular, Cellular, and Translational Research grant P30 GM106394 and Center for Molecular, Cellular and Translational Immunological Research grant P30 GM103415. JJO was partially supported by NIH R01 AI139133 grant. MEK and RMT were supported by Immunology Training Program (NIH/NIAID T32 AI007363). BSR was supported by the Dartmouth Cystic Fibrosis Training Program (NIH/NHLBI T32 HL134598). RAC, LAL, and JES were partially supported by NIH R01 AI130128. JS is a CIFAR Fellow in the Fungal Kingdom: Threats and Opportunities program. CHK was partially supported by NIH F31 AI138354. The funders had no role in the preparation or publication of the manuscript.

---

**SUPPLEMENTAL TABLE 1.** Strains used in this study. The strain identifiers, genotypes, and source information for all fungal strains included in this study.

| <u>Strain ID</u> | <u>Genotype</u> | <u>Source</u> |
| --- | --- | --- |
| AF293 | Reference Strain | Nierman *et al.* 2005<sup>68</sup> |
| CEA10 | Reference Strain | Girardin *et al.* 1993<sup>69</sup> |
| LH-EVOL | Experimentally evolved from AF293 | This study |
| AF293 Δ*sskA* | AF293 mutant lacking *sskA* | This study |
| AF293 Δ*sskA*<sup>RC</sup> | Ectopic complementation strain of the AF293 Δ*sskA* mutant | This study |
| AF293 Δ*sakA* | AF293 mutant lacking *sakA* | Xue *et al.* 2004<sup>13</sup> |
| AF293 Δ*mpkC* | AF293 mutant lacking *mpkA* | Reyes *et al.* 2006<sup>52</sup> |

**SUPPLEMENTAL TABLE 2.**
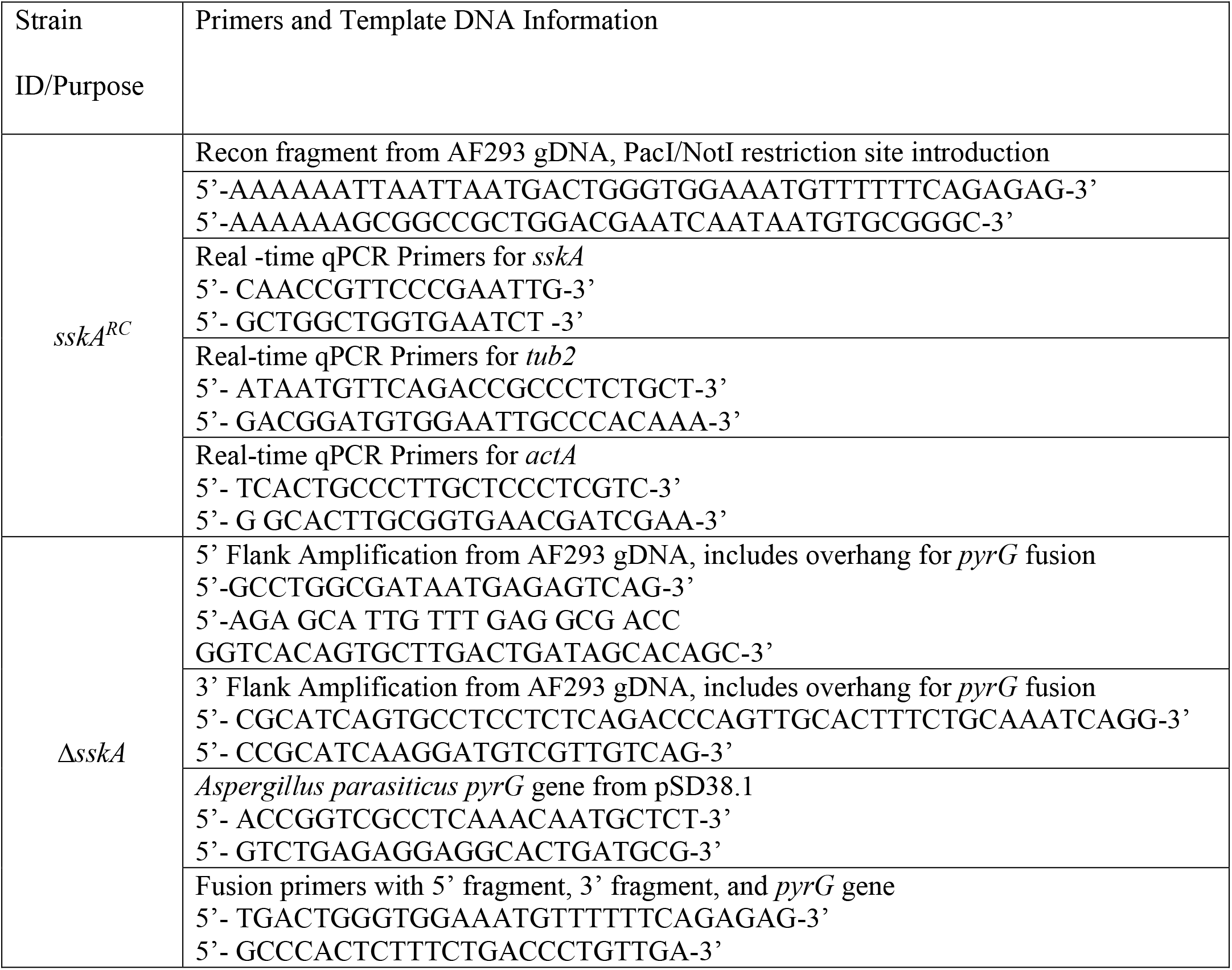
Primers used to generate AF293 Δ*sskA* and AF293 Δ*sskA::sskA*.

**Supplemental Table 3.**
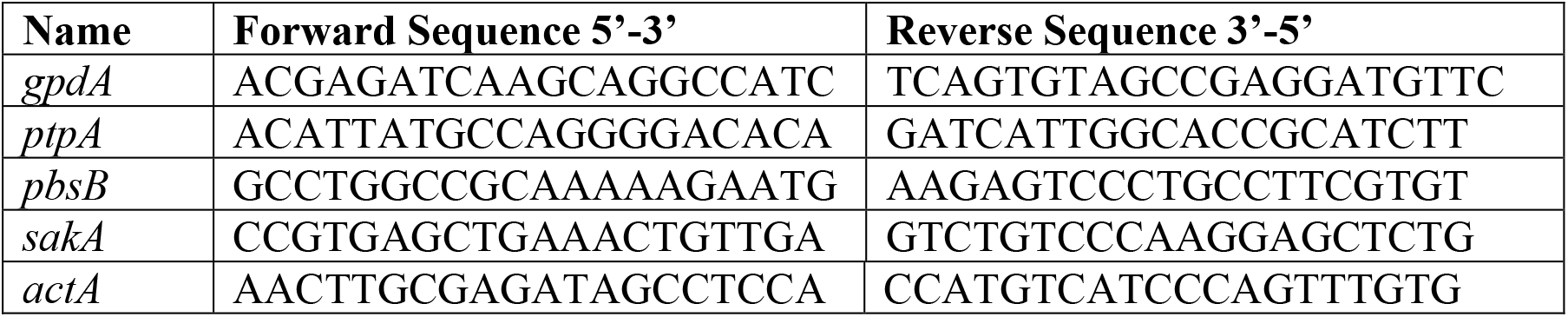
Primers used for transcriptional analysis of SakA-dependent genes.

**Supplemental Table 4.** Protein Sequences used in BLASTp analysis in non-redundant database, GenBank Accession.

|  | **Af293** | **CEA10** | **Similarity (%)** |
| --- | --- | --- | --- |
| **TcsB** | XP_001481640.1 | EDP53731.1 | 97.87 |
| **YpdA** | XP_751798.2 | EDP50401.1 | 100 |
| **SskA** | XP_753797.1 | EDP51583.1 | 100 |
| **SskB** | XP_752459.1 | EDP56327.1 | 100 |
| **PbsB** | XP_752961.1 | EDP56828.1 | 100 |
| **SakA** | XP_752664.1 | EDP56531.1 | 100 |
| **MpkC** | XP_753727.2 | EDP51653.1 | 98.94 |

**Supplemental Figure 1.**
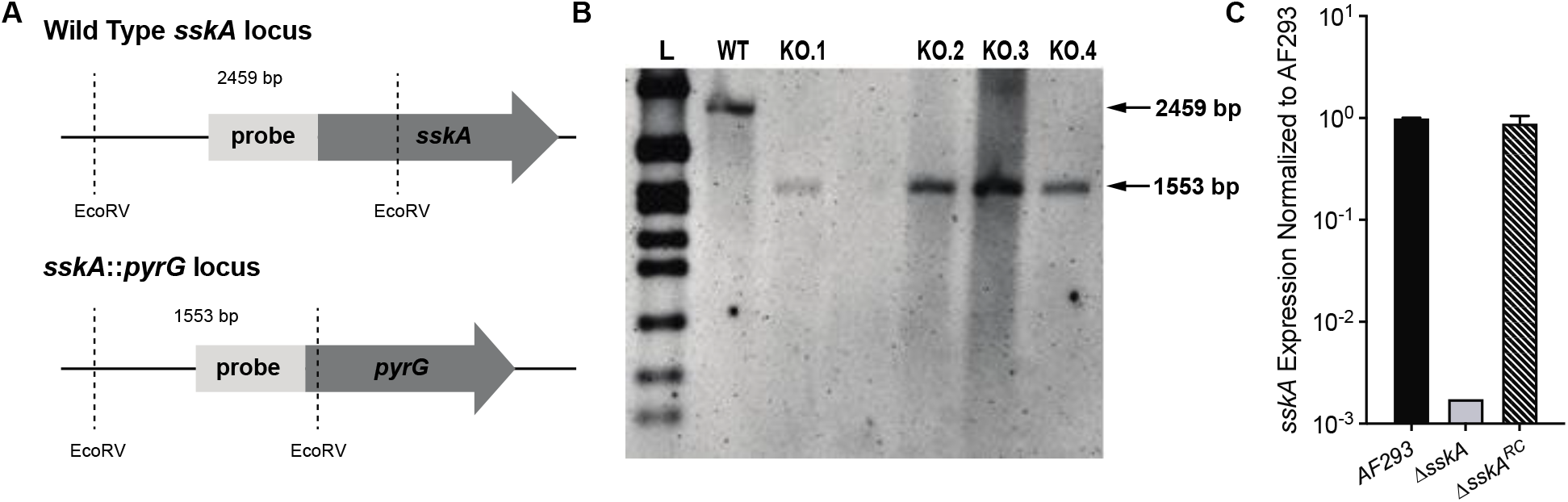
Construction and confirmation of *A. fumigatus* Δ*sskA* and Δ*sskA*^RC^ strains. (A) Schematic of *sskA* gene replacement in *A. fumigatus* strain AF293.1 with expected band sizes following genomic DNA digestion with EcoRV. (B) Southern blot analysis of EcoRV-digested genomic DNA with a 1088 bp digoxigenin-labeled probe reveal successful gene replacement of *sskA* with a single copy of *pyrG* in multiple transformants (KO.1, KO.2, KO.3, KO.4). (C) Real-time quantitative PCR results for *sskA* mRNA levels in the WT (AF293), Δ*sskA* (KO.4), and *sskA*^RC^ strains from 18 hour liquid cultures.

**Supplemental Figure 2.**
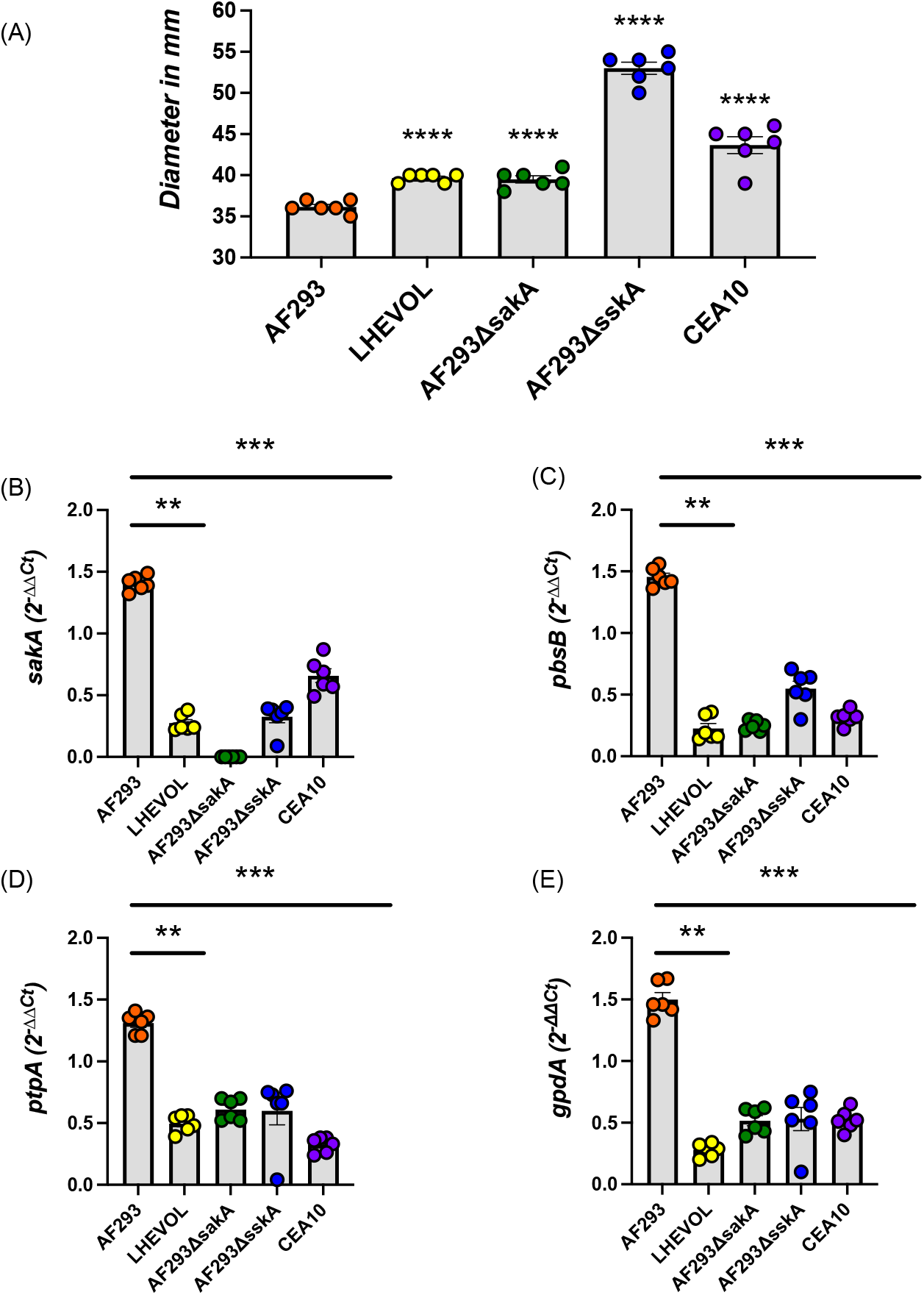
CEA10 and AF293 Δ*sskA* strain demonstrates deficiencies in *sakA*-dependent osmotic stress response. (A) Diameter of colonies grown on GMM for 72 hours in millimeters (mm). (B-E) A 30-hour biofilm was transferred to fresh liquid GMM media supplemented with 1M NaCl for 30 minutes at 37°C. Quantitative RT-PCR was used to measure the change in transcript production of (B) *sakA* and *sakA*-dependent genes (C) *pbsB*,(D) *gpdA*, and *ptpA*. Ct values were initially adjusted to *actA* for delta Ct, then 1M NaCl was compared to GMM for delta delta Ct values. Data are pooled of results from 2 independent experiments. Statistical significance was assessed using (B) a one-way ANOVA with Tukey’s test and (D) Mann-Whitney test and one-way ANOVA with a Kruskal-Wallis test. (**) indicates a p-value ≤ 0.01; (***) indicates a p-value ≤ 0.001; (****) indicates a p-value ≤ 0.0001.

**Supplemental Figure 3.**
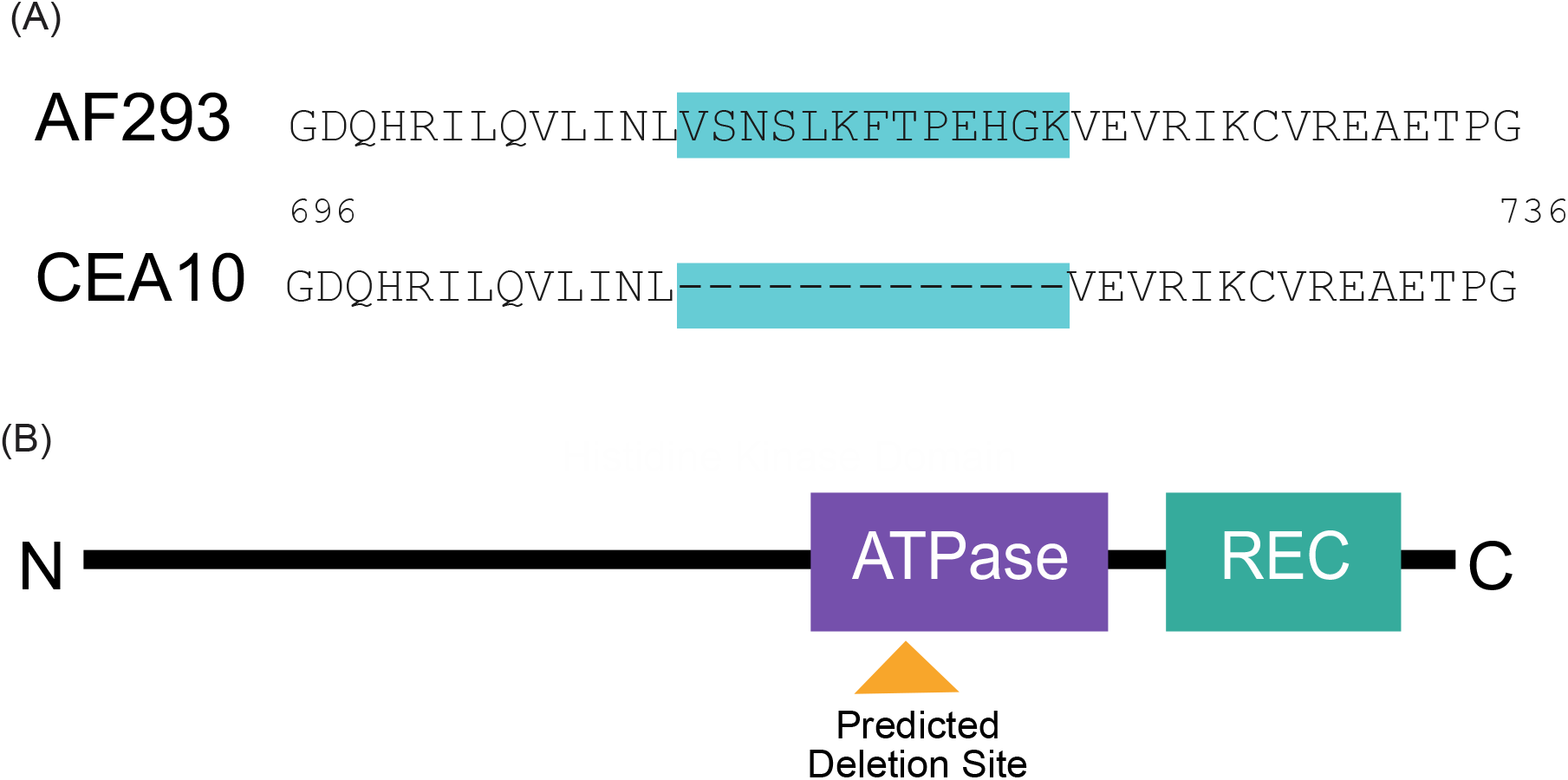
Amino acid alignment of sensory histidine kinase response regulator TcsB in AF293 and CEA10. (A) Amino acid sequence demonstrating deletion (709-721, XP_001481640.1) within the predicted ATPase (696-863, XP_001481640.1) in the histidine kinase domain in CEA10 when compared to AF293. (B) Schematic of TcsB protein demonstrating relative position of amino acid deletion in CEA10.

